# Trade-off Between Resistance and Persistence in High Cell Density *Escherichia Coli* Cultures

**DOI:** 10.1101/2024.01.29.575348

**Authors:** F. Beulig, J. Bafna-Rührer, P.E. Jensen, S.H. Kim, A. Patel, V. Kandasamy, C. S. Steffen, K. Decker, D.C. Zielinski, L. Yang, E. Özdemir, S. Sudarsan, B.O. Palsson

## Abstract

Microbes experience high cell density in many environments that come with diverse resource limitations and stresses. However, high density physiology remains poorly understood. We utilized well-controlled culturing systems to grow wild-type and metabolically engineered *Escherichia coli* strains into high cell densities (50–80 g C**_dry cell weight_** L^-1^) and determine the associated transcriptional dynamics. Knowledge-enriched machine-learning-based analytics reveal distinct stress-related gene expression patterns that are consistent with a fundamental trade-off between resistance and persistence. We suggest that this trade-off explains observed growth arrests in high-density cultures and that it results from the disruption of cellular homeostasis, due to reallocation of limited cellular resources from resistance functions towards maintenance requirements of engineered production pathways. This study deepens our understanding of high-density physiology and demonstrates its importance to fundamental biomanufacturing challenges.

## Introduction

A comprehensive understanding of high cell density microbial physiology has remained elusive despite broad and fundamental importance, such as in the context of infectious diseases and industrial biotechnology. For example, achieving high cell densities is key to economic viability of many biomanufacturing processes that are being developed to support sustainable lifestyles^1,2^. Unfortunately, detailed experimental interrogation of microbial physiology at high cell densities has been hampered by the lack of suitable methodologies but is now enabled by advancements in “omics” approaches and the development of highly controlled, parallelized bioreactors^2^.

Dense microbial populations face various sources of stresses that, among others, include transient shortage of carbon and nutrients and accumulation of inhibitory or toxic metabolites^3–7^. Stresses need to be balanced with other vital cellular processes to maintain viability. Due to the likelihood of encountering a linked set of adverse challenges, microorganisms are equipped with a global stress response that confers resistance against a variety of stresses, even those not yet encountered or seemingly unrelated^8^. Independent of the general stress response, some stress response pathways enable cells to address individual stressors by selectively activating specific gene sets^9,10^. However, maintaining resistance comes at great metabolic cost and the upregulation of stress response genes typically results in a concurrent downregulation of genes involved in growth-associated processes^9–11^. Indeed, “resistant” phenotypes have been shown to be outcompeted by “sensitive” phenotypes that do not incur the maintenance cost of constitutive stress readiness^12^. If resistance functions are exhausted, persistence remains as an alternative survival strategy. Unlike resistance, persistence is a transient physiological state that is characterized by a mixture of growing and growth-arrested cells that result from bistable expression states within a population, allowing cells to adapt and resume growth after an otherwise lethal stress exposure^13,14^.

*Escherichia coli*, an extensively studied and widely used workhorse in biotechnology, can achieve >100 g C_dry cell weight_ L^-1^ of cell density^3,4^. The natural biosynthetic capabilities of *E. coli* have been expanded by introducing heterologous pathways to develop new biosynthetic pathways. However, engineered pathways may disturb the intracellular processes and resource distribution that have been optimized by natural evolution^15,16^. The associated metabolic burden reduces cellular fitness, as limited cellular resources must be reallocated to maintain physiological and biochemical homeostasis^16,17^. Thus, cellular productivity remains difficult to predict and growth arrests can be observed, especially at high cell densities.

It remains uncertain in what way dense populations of *E. coli* balance resistance and proliferation functions. The primary objective of this study was to investigate transcriptomic responses associated with high cell density, especially under the metabolic burden of engineered production pathways. We grew different *E. coli* strains in well-controlled bioreactor environments to high cell densities and utilized knowledge-enriched, machine-learning-based analysis to facilitate interpretation of complex transcriptomic responses. Our findings suggest that, at high cell densities, the degree of metabolic burden imposed by engineered production pathways modulates previously unrecognized growth-versus-survival transitions.

## Results and Discussion

### Maintenance requirements distinguish growth of E. coli wild type and engineered strains into high cell density

High cell density cultivation experiments were conducted following a common two-stage, batch-to-fed-batch, fermentation strategy in parallel sets of bioreactors (Fig. 1A). We studied five groups of *E. coli* strains with varying degree of engineering (Fig. 1B; see Materials and Methods for details), including a wildtype BW25113 control strain (WT), different single gene knockout strains (SGKO), two tryptophan production strains (HMP3071 and empty plasmid control SDT551), and a plasmid-carrying melatonin production strain (HMP3427). After initiation of the fed-batch phase, growth was maintained with exponential feeding strategies, providing a continuous supply of media, O_2_ and pH buffering capacity to all strains (see Materials and Methods for details). We inferred four major fermentation phases (I–IV; Fig. 1A), corresponding to the transition from batch to fed-batch mode (I/II, 0 h), changes in melatonin and tryptophan formation (II/III, 10 h), and growth arrest of HM3427 (III/IV, 20 h).

**Figure 1.**
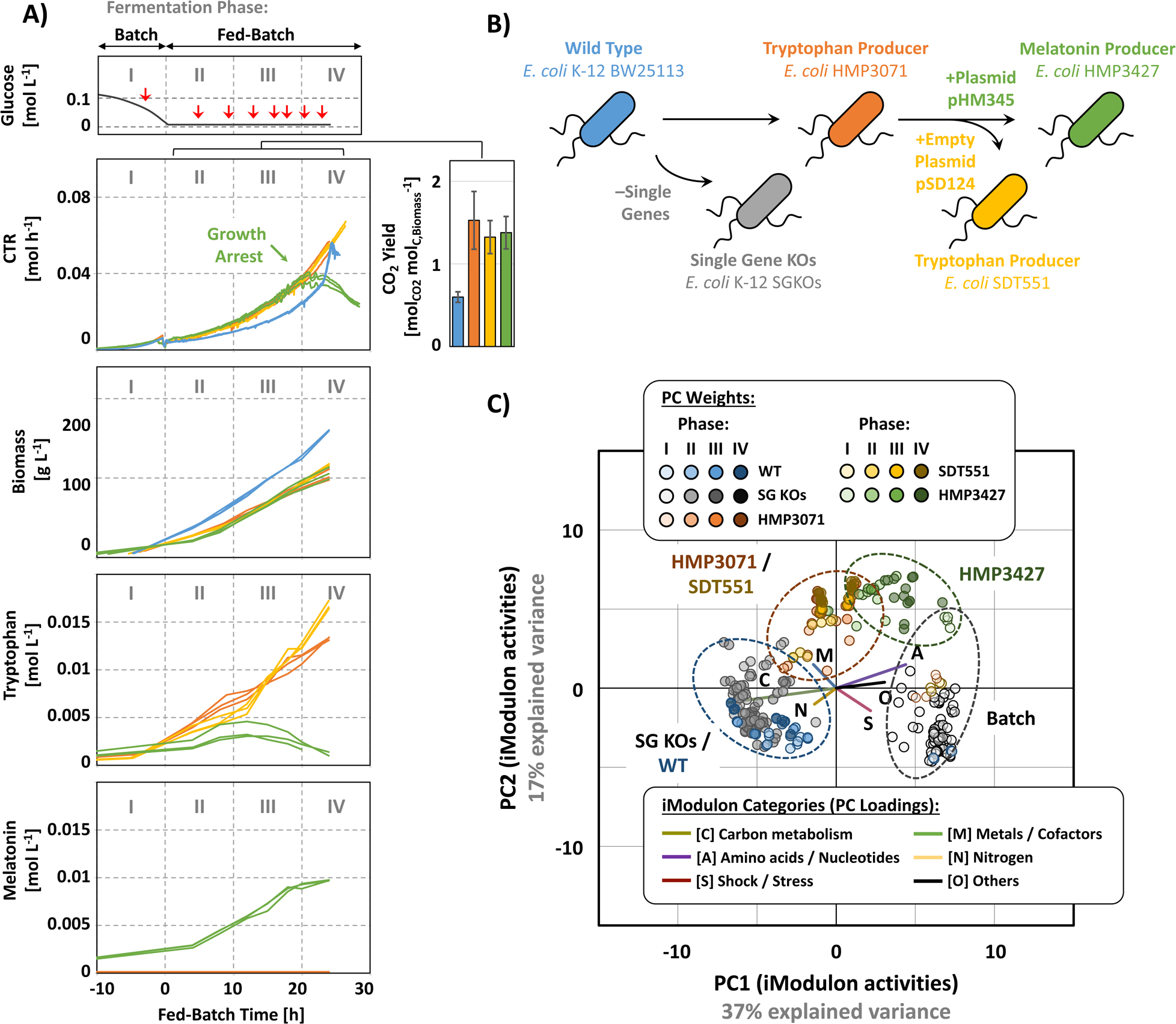
Integrated process and transcriptome diagnostics of high cell density fermentations. (A) Representative time profiles of fermentations from batch (phase I) to fed-batch (phases II–IV) with *E. coli* WT reference strain (blue), as well genome-engineered production strains HMP3071 (orange), SDT551 (yellow) and HMP3427 (green). Time points for RNA-Seq sampling are indicated by red arrows. Fermentation phases I–IV were chosen according to the transition from batch to fed-batch mode (0 h), changes in product formation (10 h), and beginning of the HM3427 growth arrest (20 h). (B) Schematic outline of strain lineage. (C) Biplot of principal component 1 (PC1) and 2 (PC2) from transcriptome samples at different stages of the fermentation.

Over the time course of cultivation, the increase in cell density was most rapid for the WT strain, with maximum biomass concentrations (average±standard deviation) of 80±3 g_Cell Dry Weight_ L^-1^ for WT (N=6) and 51±4 g_Cell Dry Weight_ L^-1^ for production strains (N=18). In contrast, CO_2_ production of WT strains was about 30% lower than production strains. Melatonin and/or tryptophan production accounted for <1% of the carbon from oxidized glucose in production strains. Thus, as evidence for the metabolic burden imposed by the engineered pathways, all genome-engineered strains sustained maintenance requirements by allocating almost twice as much of the carbon and energy from glucose oxidation towards catabolism (CO_2_ yield per formed biomass; Fig. 1A). Consistent with a growth arrest towards the end of the fed batch fermentation, CO_2_ formation of HMP3427 plateaued at the end of phase III at 0.04 mol_CO2_ h^-1^ and, following this trend, both melatonin and tryptophan production substantially decreased.

### Contrasting transcriptome structures of E. coli wild type and production strains at low and high cell density

To resolve the transcriptome dynamics of WT and production strains from low to high cell densities, we obtained at least eight replicated transcriptome samples over the course of each fermentation (Fig. 1A). Expression of heterologous genes in production strain HMP3427 accounted for 45±10% of total mRNA reads (average±standard deviation; N=32), thus occupying a significant fraction of the transcriptome. For an initial exploration of differences in gene expression patterns among the strains, we reduced the dimensionality of all transcriptomic datasets with Principal Component Analysis (Fig. 1C). The first two principal components, together explaining 54% of expression variability, revealed consistent, condition- and strain-specific gene expression patterns that resulted in clusters representative for batch phase samples of all strains (low cell density) or fed-batch samples of individual strains (high cell density). One of the fed-batch cluster was shared by samples of strains HMP3071 and SDT551, highlighting that both strains share similar global transcriptome dynamics. SGKO strains shared a cluster with the WT strain. Overall, the transition from batch to fed-batch mode had the most impact on the global transcriptome structure of each strain, and each strain appeared to respond differently to the transition.

Since the insight into transcriptomic changes using PCA was coarse-grained, we performed Independent Component Analysis (ICA) of the transcriptomic dataset (see Materials and Methods for more details). ICA allows for detailed knowledge-enriched insight into transcriptome dynamics as represented by iModulons, i.e., unique sets of independently modulated genes that reflect the transcriptional regulatory network^11^. By reducing the high-dimensional nature of transcriptome data into a smaller set of biologically meaningful gene expression patterns, iModulons facilitate interpretation of complex transcriptomic responses. Among all analyzed strains and fermentation phases, iModulons captured >90% of genes that were significantly upregulated (179 ±34; Δlog tpm >2, FDR <0.1) or downregulated (77 ±22; Δlog tpm <–2, FDR <0.1). Out of 194 iModulons, 48 were considered putatively high-density specific with >5 difference in activity between the fed-batch phase and the batch phase (|ΔA| >5, FDR <0.1). Between the different media compositions, “High Fe/Zn” and “Low Fe/Zn”, only Zinc, FliA and FlhDC iModulons were highly active (|ΔA| >5, FDR <0.1), suggesting an essential role of zinc in the expression of genes involved in flagellar synthesis^18^. Consistent with PCA-based clustering (Fig. 1C), a comparison of fed batch samples of HMP3071 against SDT551 did not reveal activated iModulons (|ΔA| >5, FDR <0.1). A comparison of SGKO against WT samples revealed an activation of sequence insertion elements (IS1 iModulon). As will be discussed in the following sections, several of the putative high-density-specific iModulons showed systematic variability over the course of the fermentation, revealing underlying regulatory mechanisms. We could not identify a pronounced correlation of iModulons with OD (*Spearman* |R^2^| >0.7, p <0.05), possibly due to the absence of universal, continuous relationships of iModulon activities with cell density itself. Differential iModulon activity during growth phases I-IV thus reflected transcriptome re-allocation with increasing OD.

To reveal functional associations and underlying regulatory relationships that govern stress-related responses at high cell densities, we searched the transcriptome datasets of all investigated strains for highly significant correlations among the 48 activated iModulons (|*Spearman* R^2^| >0.7, p <0.05; Fig. 2A). Such correlations have been used to identify stimulons^11,19^. We detected three major clusters, represented by (i) RpoS and GadXW (resistance-related), (ii) Crp and FlhDC (carbon metabolism- and motility-related), (iii) MarA and SoxS (chemical stress-related) as their largest and most connected iModulons, respectively. The broad variability of detected iModulon activity changes allowed us to robustly separate the transcriptomic state of individual strains, as it captured a substantial portion of gene expression states in the background data set (>1000 RNAseq profilies; PRECISE-1K)^19^. Further, deviation of iModulon–iModulon activity correlations from the background dataset, generated from low density cultures, highlights both strain- and high cell density-specific gene regulatory dynamics. In the following sections we will provide a systems-level description of how coordinated iModulon activities represent well-defined underlying molecular processes that reflect a novel trade-off between resistance- and persistence-like stress responses governing high cell density physiology.

**Figure 2.**
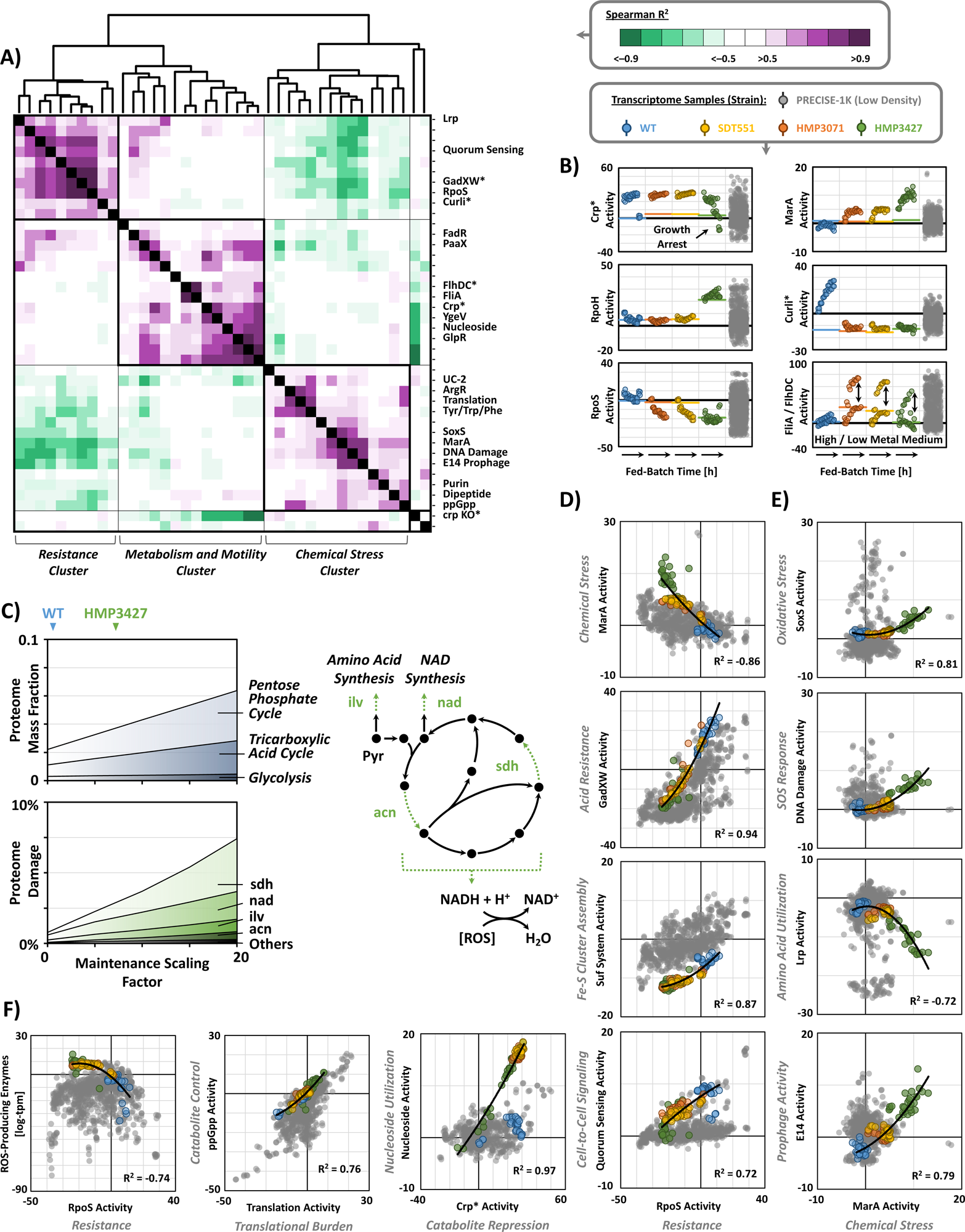
iModulon analysis reveals trade-off between resistance and persistence functions. (A) *Spearman* correlation matrix for activity levels of high cell density-specific iModulons with significantly higher or lower activity in fed-batch phase as compared to the batch phase (|A| >5, FDR <0.1). iModulons with size (number of genes) and/or connections (*Spearman* |R^2^| >0.7) >10 are highlighted. Clusters representing stimulons were named according to iModulons with largest size and most connections. (B) Activity of major iModulons Crp, RpoH, RpoS MarA, Curli, and FliA / FlhDC over fed-batch time. Horizontal lines represent average activity during the batch phase. Vertical arrows indicate FliA / FlhDC activity difference between cultivations with low and high metal supplementation, suggesting the role of Zn as an essential motility cofactor. (C) Proteome allocation for central carbon metabolism and proteome damaging reactions, as calculated with the OxidizeME model constrained by process and RNA-seq data. Predicted damage to enzymes associated with the TCA cycle are highlighted in green. (D–F) Representative iModulon correlations for global resistance (D), chemical stress (E) and general functions (F). iModulons in (A–F) denoted with a * represent the sum of iModulons with the same annotation but differing suffix (*i.e.* Crp*, crp KO*, Curli*, FlhDC*, and GadXW* represent the sum of Crp-1/2, crp KO-1/2, Curli-1/2/3, FlhDC-1/2, and GadXW-1/2, respectively).

### Carbon and energy limitation promote chemical stress at high cell density

The transition from low to high cell densities is inevitably accompanied by a transition from unrestricted growth (batch phase) to growth under substrate-limitation (fed-batch phase) and a necessity to rebalance catabolism and anabolism to support heightened maintenance requirements and biosynthetic needs^20,21^. *E. coli* responds to carbon- and energy limitation by synthesizing cyclic adenosine monophosphate, cAMP, and its binding to the cAMP receptor protein, Crp^20^. Thus, at the start of the fed batch phase, all strains upregulated multiple cAMP-Crp-regulated iModulons (crp KO-1/2, Crp-1/2, FucR/ExuR, GlcC, GlpR, MalT, Nucleoside, PaaX, Propionate, PTS II, RhaT), containing more than 200 genes that allow catabolic flexibility under carbon starvation (Metabolism and Motility Cluster; Fig. 2A and 2B). Consistent with the detected growth arrest of HMP3427, activity of all cAMP-Crp-regulated iModulons diminished towards the end of the fermentation.

High-density cultures must produce a significant amount of energy to meet their metabolic demands, which result in continuous formation of toxic radicals, as about 0.5% of consumed oxygen inevitably turns into reactive oxygen species (ROS) that cause damage to vital macromolecules, such as DNA, RNA, proteins, lipids^22,23^. To characterize the impact of ROS on WT and production strain metabolism *in silico*, we utilized OxidizeME, a genome-scale computational model of *E. coli* metabolism and expression, that incorporates a multiscale description of ROS stress response and proteome damage^24^. Consistent with higher catabolic fluxes in production strains (Fig. 1A), our model predicts a 6-fold increase in maintenance cost over the WT strain and the associated demand for ATP and reducing equivalents is supported by an enhanced flux into pentose phosphate cycle and tricarboxylic acid cycle (Fig. 2C). However, higher catabolic fluxes also lead to a 3-fold increased generation of ROS and associated damage to the proteome. If the calculated intracellular production of ROS (∼90 µmol L_Cell_^-1^ s^-1^) outstrips the cell’s NADH-dependent capacity for detoxification and ATP-dependent repair capacity, e.g. due to substrate limitation, intracellular ROS accumulation would breach toxic threshold and trigger growth arrest (>0.5 µmol L_Cell_^-1^)^22^.

### Knowledge-enriched transcriptome analysis reveals trade-off between resistance and persistence

In *E. coli*, the general stress resistance mediator is σ^38^, RpoS, a sigma subunit of RNA polymerase that partially replaces house-keeping sigma factor σ^70^, RpoD, to regulate more than 500 genes that allow rapid responses to threatening environmental conditions^8,10,25^. We identified an upregulation of the RpoS iModulon in the WT strain and distinct positive correlations with multiple stress- and resistance-related iModulons, including GadXW, Suf System, RpoE, NRZ and Quorum Sensing (*Spearman* R^2^ >0.7), that are consistent with a general stress readiness (Resistance Cluster; Fig 2A, 2B and 2D). For example, acid stress is one of the most common environmental stresses for *E. coli*, and, although the cultivation was performed under controlled pH conditions, the WT strain activated GadXW iModulons containing central components of the glutamate decarboxylase acid tolerance system^8^. The WT strain also coordinated the downregulation of ROS-producing enzymes (Fig. 2F), an upregulation of antioxidant systems as part of the RpoS iModulon, including catalases KatE, KatG and DNA-binding ferritin Dps, as well as the Suf System for reactive Fe-S cluster assembly^22,26^.

RpoS and associated stress- and/or resistance-related iModulons were significantly downregulated in production strains (Fig. 2B and D), and, as part of a distinct negative correlation (*Spearman* R^2^ = -0.86), we identified the concurrent upregulation of chemical stress-related iModulons MarA and SoxS (Chemical Stress Cluster; Fig. 2E). In fact, MarA activities in HMP3427 exceeded those in almost the entire low density background dataset. Together with SoxS, MarA acts as transcriptional regulator of overlapping regulons that contain genes involved in stress response to inhibitory and toxic chemical species, including antibiotics and redox cycling compounds, such as ROS^27^. As evidence of the cells attempt to shield themselves from harmful chemicals, we found an upregulation of multidrug exporters MdlAB and AcrAB, as part of the SoxS iModulon, concurrent to a downregulation of porins and diverse uptake systems in the ROS TALE and Lrp iModulons that is mediated by non-coding RNA micF under regulatory control of SoxS (Fig. 2E)^10,27^.

In the absence of resistance or tolerance functions, severe stress signals may trigger persistence and associated growth arrest as an alternative survival strategy^13,14,28,29^. At its core, persistence is a phenotypic response to strong perturbation of metabolic homeostasis, involving multifactorial and redundant molecular mechanisms^13,30^. The most significant persistence triggering mechanisms involve sensitization through suppression of global resistance (RpoS response), as well as stress responses towards starvation (nutrient and ATP limitation), disruption of proteostasis (heat shock response), DNA damage (SOS response), and inhibitory or toxic chemicals (multidrug response)^13,14,28–36^. As an emergent property of cell-to-cell communication in dense microbial populations, persistence has been extensively studied in the context of infectious diseases and antibiotic resistance^14,29,37,38^. However, persistence has not been reported to be of relevance in bioprocess systems so far. We found recurring evidence for activation of persistence-mediating mechanisms in the investigated production strains, and almost exclusively at high cell densities, as described in the following sections (Fig. 2A-E; Fig. 3).

**Figure 3.**
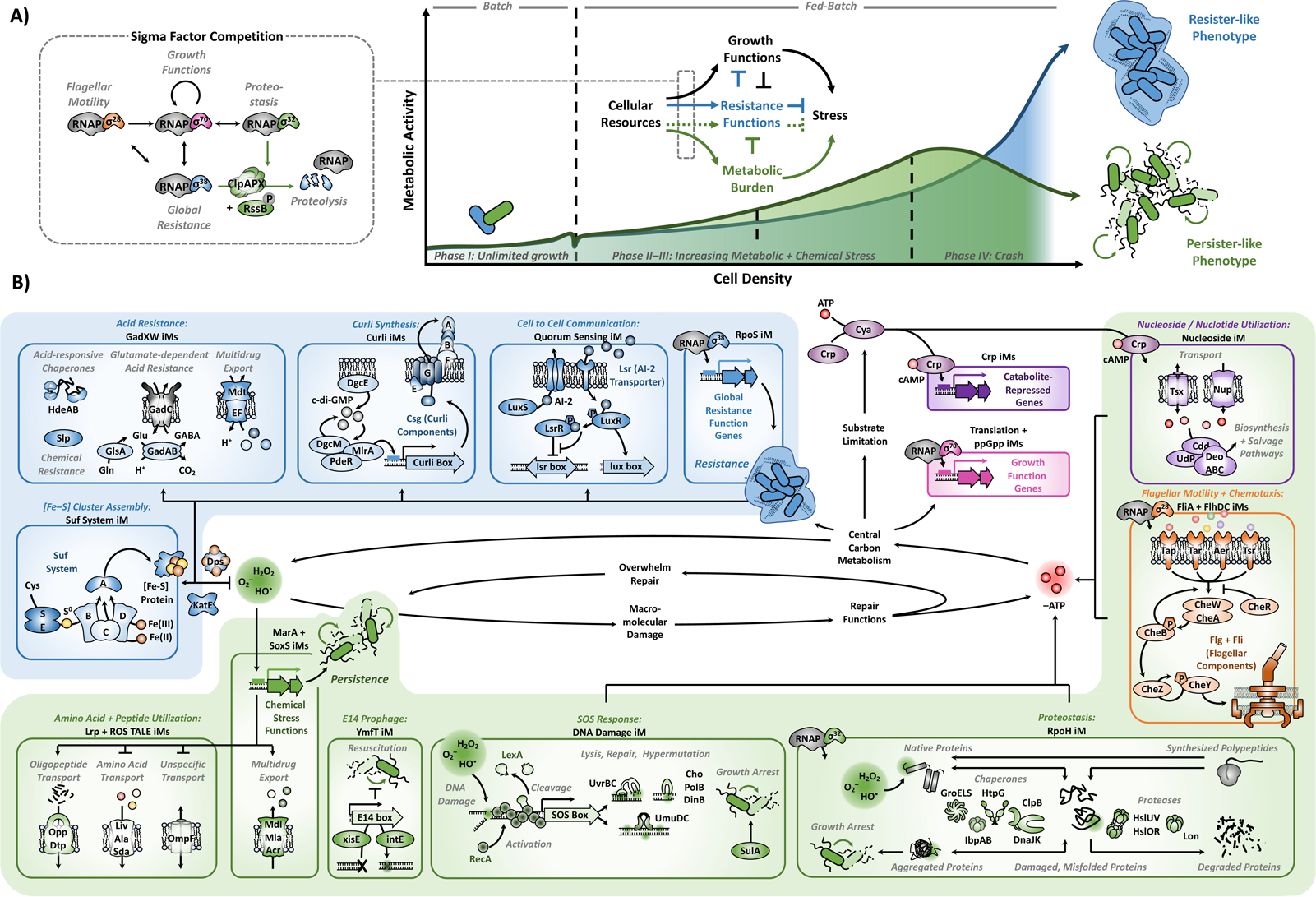
Interaction of cellular processes that are independently regulated during growth to high cell densities. Graphical summary of (A) the sigma factor regulatory network, controlling the distribution of cellular resources, and (B) the molecular processes, represented by activated iModulons, shaping the resistance-versus-persistence trade-off at high cell densities. Blue arrows and boxes represent processes linked to resistance functions in the wild type strain “WT”. Green arrows and boxes represent processes linked to persistence in the production strain “HMP3427”.

### Genotoxic and proteotoxic stress modulate persistence at high cell density

Unmitigated exposure to oxidative stress appeared to trigger SOS responses resulting in continuously increasing activity of the DNA Damage iModulon over the course of the fermentation, especially in production strain HMP3427 (Fig. 2E). The SOS response modulates bacterial persistence by acting as a signaling pathway that complements other stress response and providing DNA repair functions that are essential for transitioning back from dormancy to a metabolically active state^36^. Most of the genes belonging to the SOS regulon are repressed by LexA and are induced when LexA is self-cleaved upon association with RecA that is activated by binding to single-stranded DNA resulting from DNA damage^39^. According to the affinity of LexA for SOS genes, we found that HMP3427 not only activated “weaker” SOS functions, such as UvrAB-dependent DNA repair, but also “stronger” SOS functions, such as UmuDC-dependent mutagenesis repair and SulA-dependent growth arrest^40^. Further, we found increasing activation of the YmfT iModulon, containing E14 cryptic lambdoid prophage elements (Fig. 2E). Under stress, cryptic prophages have been shown to provide diverse benefits for survival and E14 is among the most commonly activated, especially during the SOS response^41^. Although prophages do not appear to influence persister formation they inhibit their transition back to a metabolically active state^42^.

Consistent with challenges to maintain proteostasis during heterologous protein expression, we found exceptionally high activity of the RpoH iModulon for HMP3427 cells during all four phases of the fermentation (Fig. 2B). The RpoH iModulon contains multiple chaperones (DnaJK, ClpB, IbpAB, GroESL, GrpE, HtpG, HslOR) and proteases (Lon, HslUV, ClpXP) that act as translational control systems. An upregulation of RpoH can be accompanied by downregulation of RpoS, likely because of direct proteolysis, in an effort to redirect cellular resources to support the energy-intensive functionality of ATP-dependent chaperones and proteases^43,44^. Perturbation of protein homeostasis by stress and low levels of intracellular ATP favors the formation of protein aggregates, which, apart from their toxicity, facilitate persister cell formation and growth arrest^31,45^. We found ibpA and ibpB among the highest expressed genes in HMP3427 over all phases of the fermentation (11±0.5 and 11.5±0.3 log-transformed transcripts per million, respectively). These chaperones are reported to be exclusively upregulated during extensive protein aggregation, to attenuate the cell’s global stress response, and to impede protein disaggregation^31,46^. The low metabolic flux associated with growth arrest impedes the ATP-dependent clearance of protein aggregates that is necessary for resuscitation, possibly creating a feedback loop that eventually results in cell death. Indeed, an increasing upregulation of oligopeptide transport systems by production strains, including symporters DtpD and DtpA, as part of the Lrp and SoxS iModulon, and high-affinity ABC transport system Opp, as part of the ROS TALE iModulon, points towards recycling of exogenous proteins from lysed cells that intensifies with growing cell densities (Fig. 2A and E).

### Impact of high cell density on cell-cell communication, motility, and biofilm formation

During substrate limitation, *E. coli* adopts ATP-mediated intercellular communication that results in pronounced heterogeneity of intracellular ATP concentrations, whereby accumulation of ATP in a small fraction of the population is supported by starvation of the remaining cells^47,48^. In our study, production strains strongly upregulated genes that are implicated in the utilization of extracellular ATP as part of the Nucleoside iModulon, which positively correlated with cAMP-Crp-related iModulons, indicative of substrate starvation signals (Fig. 2B). As highlighted earlier, starvation is an important trigger for persistence^33–35^. The Nucleoside iModulon includes the channel-forming protein Tsx, permeases NupC and NupG, as well as associated salvage pathway proteins Cdd, Udp, and DeoABCD. As transmission of ATP is localized, ATP-based intercellular communication is expected to become only relevant at growing cell densities. Quorum Sensing, as another form of intercellular communication, was primarily expressed in WT samples at high cell densities (Fig. 2D). The negative correlation of Quorum Sensing with MarA (*Spearman* R^2^ = -0.79) is likely established directly through repression of BhsA, as part of the MarA iModulon, and indirectly via repression of biofilm formation, distributed across the Curli iModulons^7,27^.

The trade-off between motility and biofilm formation is inversely regulated by RpoS through competition with flagellar sigma factor σ^28^, FliA, and dependence of biofilm-forming curli synthesis on the downregulation of the flagellar regulatory cascade^7^ (Fig. 2C). Consequently, an upregulation of flagellar assembly and chemotaxis, as part of the FlhDC and FliA iModulons, was restricted to production strains. Flagellar motility can mediate persistence functions. One link lies in the energy expenditure of flagellar motility, which requires extensive utilization of the TCA cycle and, in turn, leads to ATP increased production of ROS^23^. Another link lies in cell density- and communication-mediated chemical stress signaling^37^. AcrA, upregulated in production strains as part of the SoxS iModulon, was previously identified as a “necrosignal” that is released from antibiotics-killed swarmer cells to communicate an emergency state and induce chemical stress functions (MarA and SoxS iModulons) in the surviving population^37,38^.

### Molecular mechanisms modulating resource allocation at high cell densities

Microorganisms cannot maximize proliferation and stress resistance at the same time. This fundamental constraint is expressed in the “fear vs. greed” trade-off between the RpoS iModulon and RpoD-regulated iModulons that, on a molecular level, follows the competition between available sigma factor subunits for a limited number of RNA polymerase molecules (Fig. 3A). Specifically, the two growth-related iModulons ppGpp and Translation govern essential processes associated with protein synthesis and ribosomal function. Their positive correlation (*Spearman* R^2^ >0.76, p <0.05; Fig. 2F) represents the transcriptional blueprint for the allocation of limited cellular resources towards proliferation (growth or maintenance functions), highlighting a more “offensive” strategy in production strains, as opposed to a “defensive” withdrawal of precursors towards resistance functions in the WT strain. Thus, production strains prioritize outgrowing any damage accumulating resulting from a lack of resistance functions, while navigating the constraints imposed by the necessity for resource-intensive maintenance and repair functions.

### Conclusions

Taken together, our findings suggest that, at high cell densities, oxidative stress and substrate limitation renders cells under metabolic burden especially susceptible to persister cell formation and growth arrest. The integration of knowledge-enriched, machine-learning-based analysis allowed us to systematically dissect the fundamental transcriptomic structure of a trade-off between resistance- and persistence-like functions that is modulated by the coordinated activation of associated iModulons, representing well-defined molecular processes (Fig. 3A and B). Similar to the way other organisms balance resources between self-preservation and productivity (*i.e.* fight versus flight strategies), *E. coli* employs multiple layers of regulation that decide between resource-intensive investment in protection or maximizing growth at the risk of breaching proteotoxic and genotoxic thresholds. Despite the underlying regulatory complexity, positioning on the resistance-to-persistence continuum appears to be, at its core, dependent on resource allocation towards the weight of metabolic burden that follows the degree of genome engineering (WT < HMP3071 ≈ SDT551 < HMP3427). Although persistence has been intensively studied in other contexts, such as infectious diseases and antibiotic survival, its relevance in biomanufacturing environments has remained unrecognized. Consequently, our findings establish persistence and associated triggers and/or regulators as key target for the design of production strains that show resilience towards stresses of dense fermentation environments. Our study provides the first system level understanding of high cell density *E. coli* physiology and reveals complex links between transcriptome dynamics and metabolic engineering that are distinct from those at low cell densities.

## Materials and Methods

### Bacterial Strains and Cultivation Setup

Batch-to-fed-batch fermentation experiments were performed in 0.25 L Ambr^®^ fermenters (Sartorius) with 2 different media compositions (“High Metal” and “Low Metal” medium) and 4 strains, including strain WT (*E. coli* strain BW25113) and *E. coli* BW25113-derived production strains HMP3071, SDT551 and HMP3427 (genotype *folE*(T198I), *ynbB*(V197A), Δ*tnaA*, Δ*trpR*(P2–ckDDC), *obgE*(E350A), Δ*yddG*::PJ23101– sgAANAT(D63G), Δ*pfolE*::PJ23100, Δ*ldhA*::PJ23101–sgAANAT(D63G), *trpE*(S40F), Δ*fhuA*)^49^. Strain SDT551 carried empty control plasmid pSD124 (genotype Kan^R^) and strain HMP3427 carried production plasmid pHM345 (genotype J23107:*tph–pcd–asmt* Kan^R^)^49^.

The “Low Metal” batch media contained base medium (22 g L^-1^ D-(+)-glucose monohydrate, 7.6 g L^-1^ (NH_4_)_2_SO_4_, 15.7 g L^-1^ KH_2_PO_4_, 0.241 g L^-1^ MgSO_4_, 100 mg L^-1^ thiamine hydrochloride, 0.5 mL L^-1^ Antifoam 204), and 1 mL L^-1^ of trace metal solution (0.48 g L^-1^ CuSO_4_·5H_2_O, 0.54 mg L^-1^ CoCl·7H_2_O, 0.54 g L^-1^ ZnSO_4_·7H_2_O, 2 g L^-1^ CaCl_2_·2H_2_O, 41.8 g L^-1^ FeCl_3_·6H_2_O, 0.3 g L^-1^ MnSO_4_·H_2_O, 33.4 g L^-1^ Na_2_EDTA, 0.5 g L^-1^ Na_2_MoO_4_·2H_2_O). The “High Metal” batch media contained the same base medium, 3.5 mL L^-1^ trace metal solution and was additionally supplemented with 1.86 g L^-1^ citric acid monohydrate and 41.8 mg L^-1^ FeCl_3_·6H_2_O, 21.6 mg L^-1^ ZnSO_4_·5H_2_O). Initial batch fermentations were performed with base medium until complete glucose consumption, marked by a 50% decline in the rate of CO_2_ formation, triggering an automatic switch to fed-batch operation with continuous supply of the feed medium (550 g L^-1^ D-(+)-glucose monohydrate, 13.7 g L^-1^ MgSO_4_·7H_2_O), according to:

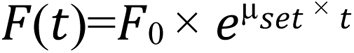

where F(t) is the continuous feed rate; F_0_ is the initial feed rate (0.535 mL h^-1^); µ_set_ is the set specific growth rate (0.11 h^-1^); and t is the process time (Lee 1996). Ambr^®^ fermenters perform online measurement of dissolved O_2_ and exhaust gas O_2_, CO_2_, as well as operating parameters. In our experiments, temperature was set at 30°C, dissolved O_2_ was controlled at 40% (cascade control with stirring speed from 1000 to 4000 revolutions per minute, air flow from 0.1 to 0.2 L min^-1^, and O_2_ flow from 0.0 to 0.05 L min^-1^), pH was maintained at 6.5 with ammonium hydroxide (4 mol L^-1^). Throughout the cultivation, samples were taken periodically by an automated liquid handler for transcriptome analysis, as well as offline analysis of biomass, tryptophan, melatonin, glucose, and organic acid concentrations.

To extend the high density transcriptome data basis for independent component analysis (described in the following), supplemental batch-to-fed-batch fermentation experiments were performed with single gene knockouts (SGKO) of *E. coli* strain BW25113 (*ΔphoB, ΔqseF, ΔbasS, ΔkdpE, ΔqseB, ΔqseC, ΔlsrR, ΔphoB, ΔqseB, ΔbaeR, ΔlsrB, ΔrpoS, ΔarcA, ΔbaeR, ΔlsrK, ΔluxS, ΔtnaA, Δcra, ΔcsgD, Δfnr, Δfur, Δnac, ΔproV, ΔprpR, ΔpurR, ΔyieP, ΔcsgD, Δfur, Δnac, ΔprpR, or ΔpurR*) in 0.25 L Ambr^®^ reactors and “Low Metal” medium, following the operational procedures described above.

### Analytical procedures

For the determination of biomass concentration during the fermentation, the optical density at a wavelength of 600 nm of the culture (OD_600_) was measured using a spectrophotometer (Jenway 7205 UV/visible; Cole-Parmer, UK). Measured OD_600_ values were converted to cell dry weight with a correlation factor of 0.31 g L^-1^ OD_600_^-1^. For the measurement of extracellular melatonin and tryptophan analysis, samples were filtered (MultiScreen® filter plates with 0.45 μm Durapore® membranes; Merck Millipore, Germany) and measured using HPLC (Dionex Ultimate 3000, Thermo Fisher Scientific, USA) equipped with a C18 column (Zorbax® Eclipse Plus C18, 4.6×100 mm, with a pre-column filter; Agilent, USA), and Diode Array Detector (DAD 3000; Thermo Fisher, USA). Separation was achieved using a gradient flow of 1 mL min^-1^ with solvent A (0.02% (v/v) acetic acid, 99.9% HPLC grade in ultrapure water) and solvent B (Acetonitrile, ≥ 99.9% HPLC grade). The following elution profile was employed: elution started with 5% solvent A, followed by three steep gradients: first to 12% from 0-1.5 min and elution was maintained until 2.5 min, second to 30% from 2.5-4.5 min and elution was maintained until 5.5 min, third to 70% from 5.5-8 min. To equilibrate the column back to its initial conditions, the gradient was decreased to 5% solvent A from 9-9.5 min and maintained until the end of run time (11 minutes). To quantify melatonin and tryptophan, 1 μL of the sample was injected, and the column temperature was kept constant at 30°C. For the analysis of glucose and short chain fatty acids (Acetate, Citrate, Formate, Lactate and Pyruvate), samples were centrifuged for 5 minutes at 16,200×g at 4°C. The resulting supernatant was diluted 1:10 in 9 mM sulfuric acid solution, filtered (MultiScreen® filter plates with 0.45 μm Durapore® membranes; Merck Millipore, Germany), and measured using HPLC (Dionex Ultimate 3000, Thermo Fisher Scientific, USA) equipped with an ion exclusion column (Aminex® HPX-87X, 300×7.8 mm; BioRad, USA), as well as refractive index and UV detectors (RI-150; Thermo Fisher, USA). Separation was achieved using 9 mM sulfuric acid solution as mobile phase with an isocratic flow of 0.7 mL min^-1^. To quantify the organic acids, 10 μL of the sample was injected, and the column temperature was kept constant at 60°C.

### RNA Sequencing and iModulon Analysis

All transcriptome samples were collected and prepared in biological duplicates, whereby either 2 ml of culture (low cell density culture, batch phase) or 0.5 ml of culture (high cell density culture, fed batch phase) was added to 6 ml of Qiagen RNA-protect Bacteria Reagent immediately after sample collection. This solution was then vortexed for 30 seconds, incubated at room temperature for 5 minutes, and then centrifuged. The supernatant was then removed and the cell pellet was stored at -80° C. The RNeasy Mini Kit (Qiagen) was used to extract and purify RNA from the cell pellets per vendor protocol, including on-columns RNase-Free DNase treatment (Qiagen) for 30 minutes at room temperature. RNA was quantified using qubit fluorometer (Thermofisher) and integrity was assessed using fragment analyzer (Agilent Technologies). Ribosomal RNA was removed using QIAseq FastSelect –5S/16S/23S kit (Qiagen). Sequencing libraries were prepared with the KAPA Stranded RNA-Seq Library Preparation Kit (Kapa Biosystems) and assessed using fragment analyzer (Agilent Technologies). The libraries were then subjected to paired end sequencing on Illumina Nextseq using Nextseq Mid Output kit (Illumina) with a read length of 150 bp. The raw sequencing reads were demultiplexed based on sample indexes using basespace (Illumina) and the fastQ files were used for further processing, as described previously (github.com/avsastry/modulome-workflow)^11^. The resulting, high-quality RNA-sequencing datasets from this study were combined with available, low density PRECISE-1K datasets (github.com/SBRG/precise1k) to compute iModulons by performing independent component analysis, as described previously^11,19^. Briefly, 100 iterations of the OptICA algorithm were performed across a range dimensionality to find robust components which appeared in more than 50 of the iterations of an individual dimensionality. An independent component dimensionality of 400 was chosen for iModulon reconstruction, annotation, and analysis, following established protocols (github.com/SBRG/pymodulon)^11,19^. At the optimal dimensionality, the total number of non-single gene iModulons was 194. Differential iModulon activities were calculated as described previously^11^. For each iModulon, a null distribution was generated by calculating the absolute difference between each pair of biological replicates and fitting a log-normal distribution to them. For the groups being compared, their mean difference for each iModulon was compared to that iModulon’s null distribution to obtain a p value. The set of p values for all iModulons was then false discovery rate (FDR) corrected to generate q-values. For comparison between samples, a difference in iModulon activity, ΔA, >5 was considered as highly active (FDR <0.1).

### Genome-scale model of metabolism and macromolecular expression (ME-model)

We used OxidizeME, a genome-scale model of metabolism and expression (ME) with ROS damage responses^24^. Models were constrained using phenotypic data (growth rate, glucose uptake rate, oxygen uptake rate, CO_2_ production rate, organic acid production rates), and expression data, as described previously^50^. Growth was simulated over a range of maintenance costs (reaction “ATPM”) with basal intracellular ROS concentrations for the WT strain (0.2 nmol L^-1^ superoxide and 50 nmol L^-1^ hydrogen peroxide). As the steady-state level of ROS is proportionate to the rate of its formation, intracellular ROS concentrations of production strain were scaled according to the increase in oxygen uptake rate relative to the WT strain^23,24^. The percentage of the proteome allocated to glycolysis, pentose phosphate cycle and TCA cycle was calculated using the solutions from each model, according to:

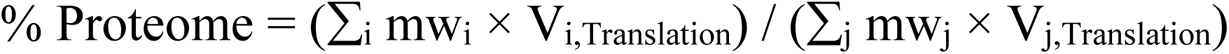

where mw_j_ and V_j,Translation_ represent the molecular weight and translation flux of the *i*-th protein in a given pathway, respectively, and mw_j_ and V_j,Translation_ represent the molecular weight and translation flux of the *j*-th protein in the entire model. Similarly, the damaged portion of the proteome was calculated, according to:

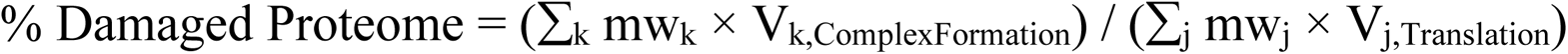

where mw_k_ and V_k,ComplexFormation_ represent the molecular weight and translation flux of all OxidizeME protein damaging reactions (denoted with the prefix “damage_”), respectively.

## Author Contributions

F.B., J.B.R., S.H.K., K.D., D.Z., L.Y., E.Ö., S.S. and B.O.P. designed the study. S.H.K. and L.Y. performed strain design and construction. F.B., J.B.R., P.E.J., S.H.K., V.K., L.Y., E.Ö., and S.S. performed experimental design and implementation. J.B.R., P.E.J., S.H.K. and V.K. performed experiments. K.D., D.Z. and E.Ö. performed initial data exploration. F.B. analyzed the data. F.B. and A.P. performed simulations. F.B. and B.O.P. wrote the manuscript with contributions from all the other co-authors.

## Data availability

RNA-seq data have been deposited to GEO and are publicly available as of the date of publication, under accession numbers GSE252784. All original data to is available at github.com/febedtu/hd_ecoli and any additional information or code required to reanalyze the data reported in this paper is available from the lead author or correspondence contact upon request.

